# *Scribble* knockdown induced metastasis tracking, identification of its associated novel molecular candidates through Proteome and *in silico* studies

**DOI:** 10.1101/2024.04.09.588727

**Authors:** Jyotsna Singh, Saripella Srikrishna

## Abstract

Metastasis is the primary cause of cancer associated death globally. Loss of function of *Scribble*, a cell polarity regulator/tumor suppressor gene, is associated with many forms of human cancer but its role in cell proliferation and metastasis remains unknown. We generated metastatic cancer in *Drosophila* using UAS-GAL4 system, through knockdown of *Scribble* in the wing imaginal discs and tracked metastasis events from 0hr early pupae to 84hrs late pupae using fluorescence microscope. Here, we report, for the first time, that knockdown of *Scribble* alone could lead to the development of primary tumor in the wing imaginal discs, which is capable of establishing metastasis, leading to secondary tumor formation, eventually resulting in absolute pupal lethality. Further, we checked a metastasis biomarker, MMP1 levels during pre-and post-metastatic phases in *Drosophila* pupae using qRT-PCR and Western blot analysis. In addition, we analyzed the proteome of *Scribble* knockdown induced tumor-bearing pupae by 2-D gel electrophoresis followed by MALDI-TOF MS to identify novel proteins involved in the process of tumorigenesis and metastasis. We identified six differentially expressed proteins, Obp 99b, Fer2LCH,CG13492, Hsp23, Ubiquitin and Colt in *Scrib* knockdown pupae compared to wild-type and validated their expression at the transcriptional level using qRT-PCR. *In-silico* studies show these novel protein interaction with *Scrib*. Thus, our results suggested that loss of *Scrib* alone causes metastasis, without the need for cooperative interaction with oncogenic *Ras.* The newly identified Fer2LCH (ferritin) and colt proteins could be important candidates for therapeutic target against *Scrib* associated cell proliferation and metastasis.

## 1. Introduction

Cancer is a significant cause of death worldwide and as per the data stated by World Health Organization (WHO’s) International Agency for Research in Cancer (IARC) (https://gco.iarc.fr/tomorrow/), the estimated number of cancer cases increase from 19.3 million to 30.2 million by 2040. Metastasis is the one of the hallmark of cancer and 90% cases of cancer related death are due to secondary tumor formation rather than primary tumor [2].Tumor metastasis is a complex process in which cells of a malignant primary tumor migrate to new distant sites, where secondary tumors develop [3].A number of studies have shown that E-cadherin, a tumor suppressor gene, has association with EMT (Epithelial-mesenchymal transition), and plays important roles in metastasis, tumor progression and cell proliferation of cancerous cells [4]. Further, in the last few decades it has been reported that metastatic behavior of *Drosophila* tumors depends on Matrix metalloproteinases (MMPs) to support the microenvironment of cancer cells [5]. Matrix metalloproteinases (MMPs), one of the metastasis biomarker, are extracellular calcium-dependent zinc-containing proteases, which regulate many signaling pathways involved in cell differentiation, proliferation, and metastasis [6]. In humans there are 24 MMPs, of which, 7 MMPs are GPI (Glycosylphoshatidylinositol) anchored and 17 MMPs are soluble secreted proteinases [7]. *Drosophila* has only two MMPs proteases, soluble secreted protease MMP1, and membrane tethered protease MMP2, making it a simple model organism for study of the involvement of MMPs in tumor metastasis [8]. Both MMPs are able to degrade DE-cadherin mediated cell-cell junctions, MMP1 being more efficient than MMP2 [9]. This function of MMPs favors malignant cells to escape and invade other tissue upon cooperative association of polarity regulator genes with*Ras^V12^*[10]. The upregulation of oncogene, *Ras^V12^* or downregulation of cell polarity regulator gene, *Scrib* in eye imaginal disc induces non-invasive tumors whereas, the cooperative involvement of *Ras^V12^*with loss of *Scrib* gene leads to metastatic tumor induction in *Drosophila* [11]. There are a few publications available to demonstrates that MMP1 facilitates the metastasis upon loss of cell polarity regulator gene, lgl (lethal giant larve) alone [3,12]. Here we show that without cooperative involvement of any other oncogenes, the polarity regulator protein, *Scrib,* alone can favor the migration of primary tumor to invade other parts, in *Drosophila* pupae. Loss-of function mutations in *Scribble* complex genes (*Scrib, lgl and dlg*) in wing imaginal discs leads to neoplastic overgrowth and disrupted epithelial homeostasis [13,14]. Epithelial homeostasis is maintained by a set of polarity regulator genes. These includes Crumbs complex (Crumbs-Pals1-Patj-Lin-7), Par complex (Par3-Par6-aPKC) and Scribble complex (Lgl-Scribble-Dlg) localized at apical domain, subapical domain and baso-lateral domain respectively [15]. These tri-molecule complex’s or modules have tumor suppressor properties and act antagonistically to maintain apical (top) and baso-lateral (bottom) compartments of epithelial cytoskeleton [16]. During normal conditions, *Scribble* polarity regulator complex maintains epithelial tissue architecture, which is disrupted upon loss of *Scribble* in many types of cancers. Previous studies show, loss of *Scrib* or overexpression of *Ras* induce slow-growing non-metastatic benign tumor however, cooperative interaction of oncogenic *Ras* and *Scribble* gene together, they accelerated the growth of malignant tumor and ultimately leads to pupal lethality [10]. However, the mechanism by which *Scribble* planar cell polarity protein (SCRIB) promotes cell proliferation and metastasis is not yet known [17]. We investigated the effect of loss of *Scrib* alone, without collaborative association with oncogenic *Ras*, on metastasis and cell proliferation. Here, for the first time, we show that *Scrib* knockdown in cells of wing discs have sufficient potential to migrate and colonize distal parts. We tracked the metastasis upto 84hrs after pupae formation (APF) and checked the status of metastasis biomarkers, MMPs and E-Cadherin during pre and post metastatic phase in *Drosophila* tumor bearing pupae. Interestingly, we found upregulation of metastasis biomarkers upon loss of *Scrib* induced migration of cancer cells. We have further analyzed the proteome of *scrib* knockdown tumorous pupae and identified six proteins which were differentially expressed and may serve as important biomarkers involved in cancer metastasis. To our knowledge, this is the first report presenting a comparative proteomic and genomic analysis of *Scrib* knockdown tumor bearing pupae and wild-type pupae to explore the role of polarity regulator gene in cancer metastasis.

## 2. Materials and Methods

### 2.1 Chemical Reagents

In this study, the following reagents were used: Urea (SDFCL), CHAPS [3 -[(3cholamidopropyl)-dimethylammonia]-1-propane sulfonate (SRL)], DTT [Dithiothreitol (TITAN BIOTECH LTD.)], IPG Buffer (Bio-Lyte), Bromophenol Blue (SRL), Thiourea (SRL), TrisHCl [Hydrochloric Acid (BR BIOCHEM LIFE SCIENCES)], EDTA [Ethylene diamine tetra Acetic Acid Disodium salt (SRL)], PMSF [Protease Inhibitor Cocktail (SIGMA-ALDRICH)], Paper Wick (BIORAD), SDS [Sodium dodecyl Sulphate (SRL)], Glycerol (Fisher Scientific), Idoacetamide (BR BIOCHEM LIFE SCIENCES), IPG strip (BIORAD Ready Strip^TM^), Acrylamide (MP Biomedicals,LLC), APS [Ammonium Sulphate (HIMEDIA)], TEMED [N,N,N^’^, N^’^-tetramethylethylene diamine (Biochem Life Sciences), Glycine (SRL), Acetic Acid (Molychem), Tris Base (SRL), Silver nitrate (SRL), Sodium carbonate (Merck), Formaldehyde (Qualigens), Mineral Oil (BIO RAD), Bisacrylamide (Qualigens), Chloroform (Merck), Isopropylalcohol (HIMEDIA), DEPC (BR BIOCHEM LIFE SCIENCES), Agarose (Seakem^R^), RNA iso Plus (TaKaRa), 6X loading Dye (Fermentas), Random Hexamer (Applied Biosystems), dNTPs (Invitrogen), RNase Inhibitor (Applied Biosystems), Reverse Transcriptase (Ambion), SYBR (R) GREEN JUMPSTART TAQ Ready mix (Thermo Fisher Scientific).

### 2.2 Fly stocks and Culture Conditions

Wild type (Oregon-R +), UAS-*Scrib^RNAi^*, UAS-GFP, sp./CyO, dCO2/TM6B (double chromosome balancers) and wing specific Ptc-GAL4 fly strains were used for this study. Fly stocks were obtained from the Bloomington Drosophila Stock Center. All crosses were carried out at 24^0^C in standard corn-meal agar media and fly stocks were maintained in B.O.D incubator.

### 2.3 *Scrib* knockdown in wing imaginal discs

We knocked down the expression of polarity regulator *Scrib*gene using RNAi approach, specifically in monocellular epithelial layer wing imaginal discs using UAS-*Scrib^RNAi^*responder line and Ptc-GAL4 driver line (Wing imaginal disc specific GAL4). For this, Virgin female flies of UAS-*Scrib^RNAi^*were crossed to male flies of Ptc-GAL4 and the quiescent transparent white pupa of F1 generation were used for the study. Fifteen each of female and male flies were taken for the cross (Genetic cross scheme shown in fig.S1, Supplementary Information).

### 2.4 Generation of Fluorescently labeled tumors

To monitor tumor progression and metastatic behavior in living transparent pupae, we genetically labeled *Scrib* knockdown cells using wing discs specific Ptc-GAL4 with a visible marker such as green fluorescent protein (GFP; Genetic cross scheme shown in fig.S2, Supplementary Information).

### 2.5 Confocal Imaging

Wing discs from third instar larvae of Wild type and UAS-*Scrib^RNAi^* /UAS-GFP;Ptc-GAL4, were dissected out in 1X PBS respectively and fixed with 4% paraformaldehyde for 20 min followed by washing with 1X PBS (phosphate buffer saline). The tissues were mounted on glass slides with DABCO. Imaged were analyzed by Zeiss LSM-510 Meta confocal Microscope.

#### Immunostaining of larval wing imaginal discs

*Drosophila* third instar larval wing imaginal discs were dissected in 1X PBS and fixed in 4% paraformaldehyde (PFA) for 15 min at RT. The tissues were washed in 0.2% PBST (1X PBS, 0.2% Triton-X) three times, 5 min each, and blocked in blocking solution (3% bovine serum albumin (BSA) solution in 1X PBS) for 1hr at RT followed by incubation in primary antibody, a cocktail of three anti-MMP1 catD monoclonal antibodies raised against the catalytic domain (14A3D2, 3B8D12, 5H7B11 used in 1:1:1 dilution) were obtained from the Developmental Study Hybridoma Bank (DSHB), Iowa, in blocking solution for overnight at 4°C. The tissues were washed with 0.2% PBST (1X PBS, 0.2% Triton-X) three times, 5 min each, and incubated with secondary antibody anti-mouse Alexa 594 (1:1000 dilution, Cat# A-11005) in 1X PBS for 3hrs at RT and washed with 0.1% PBST (1X PBS, 0.2% Triton-X) three times, 5 min each. These tissue were further incubated with DAPI (1µg/ml in 1X PBS) for 10min then again washed with 0.1% 1X PBST (1X PBS, 0.1% Triton-X) three times, 5 min each and then samples were mounted in DABCO. Immunofluorescence images were taken using confocal microscope (Zeiss LSM-510 Meta) and processed using LSM software and arranged in Adope Photoshop.

### 2.6 Tracking of tumor progression and metastasis after puparium formation (APF) from 0 hrs to 84 hrs

The F-1 generation progenies of control pupae (UAS-GFP; Ptc-GAL4) and tumor bearing pupae (UAS-*Scrib^RNAi^*/UAS-GFP;Ptc-GAL4) were observed for tracking metastasis during pupal development. The white transparent pre-metastatic tumor bearing pupae (n=30) marked as 0 hours pupae, were collected from the sides of culture bottles and placed in glass slide sealed petri dishes lined with cotton. The petri plates were stored at 24^0^C inside BOD incubator and withdrawn for records of observations at various time points throughout pupal development upto 84 hrs APF in tumor bearing pupae verses adult wings formation in wild-type pupae under fluorescence microscope. The pupae were wetted to capture images by Nikon DS-Ri-1 high resolution camera and processed by NIS-Elements (BR) software.

### 2.7 Protein Sample Preparation

Proteins were extracted from 15 pupae each after 24 hours of pupae formation from WT and UAS-*Scrib^RNAi^* driven with Ptc-Gal4. The pupae were homogenized in homogenization buffer containing 20mM TrisHCl (pH 7.6) and PMSF (1µl/ml) cocktail, centrifuged at 10,000 rpm at 4^0^C for 20 min. and supernatants were collected in fresh 1.5ml microfuge tubes. The protein concentration was quantified by Bradford method [18].

### 2.8 Two-dimensional gel electrophoresis (2-DE)

#### 2.8.1 Isoelectric Focusing (IEF)

The first dimension, isoelectric focusing (IEF) was conducted on 11cm IPG strip of pH 3 -10 linear gradients.150µg protein was used for 2-DE. Protein sample in Lysis buffer ( 20 mMTris Buffer, 7M Urea, 2M Thiourea, 4% CHAPS, 10 mM EDTA, 1mM PMSF ) were mixed with 100µl of Rehydration buffer (7M Urea, 0.5% W/V CHAPS, 0.002% Bromophenol Blue, 0.2% w/v DTT and 0.5% IPG Buffer ) and loaded in the strip holder. The strip was rehydrated for overnight, and then set program, 200V for 1hrs, 500V for 3 hrs, 1000V of 4 hrs, 8000V for 2 hrs and 8000V for 13 hrs.

#### 2.8.2 One Dimension Electrophoresis

After IEF, sodium dodecyl sulfate-polyacrylamide gel electrophoresis (SDS-PAGE) were performed on pre cast 12% polyacrylamide gel. The IPG strip were first equilibrated in equilibration buffer 1 (6M Urea, 0.375M Tris buffer, 20% Gycerol, 2% SDS, 0.2% DTT) for 10min and further in equilibration buffer 2 (6M Urea, 0.375M Tris buffer, 20% Gycerol, 2% SDS, 2.5% idoacetamide) for 10min.The strips were transferred on to gels, which were run at 200V. Three biological replicates were used for each genotype.

### 2.9 Gel Staining

The SDS-PAGE gels were fixed in fixing solution with 20% ethanol for 20 minutes, then sensitized with sensitizing solution for 2 minutes and washed with distilled water twice. Pre-chilled AgNO_3_ (silver nitrate) solution was added and after 20 min washed with distilled water two times at 30 seconds intervals. The gel was then rinsed with developing solution for spot development with gentle hand shaking. The gels were washed twice with distilled water and termination solution was added to stop the reaction.

### 2.10 Image analysis and spot picking

Image filtration, background subtraction, spot detection, spot matching and quantitative intensity analysis were performed using PD Quest software (ver.8.0.1, Bio-Rad). After image analysis, the selected protein spot that displayed differential intensities, were excised from the gels.

### 2.11 In-gel digestion

The gel slice was diced into small pieces and placed in new microfuge tubes. The gel pieces were destained using destaining solution [1:1 Ratio of ACN (Acetonitrile) and 25mM Ammonium Bicarbonate]in 10min intervals (3-4 times) until the gel pieces become translucent. The gels were dehydrated using 100μl Acetonitrile and Speedvac till complete dryness. The gel pieces were rehydrated with 10mM DTT (100µl) and followed by incubation with Iodoacetamide (100μl) for 45min and 25mM ammonium bicarbonate solution (150μl) for 10min. Subsequently, the gel was dehydrated with 100μl Acetonitrile for thermo-mixed at 600RPM for 10min and Speedvac till complete dryness. 8μl Trypsin solution was added and incubated overnight at 37^0^C. The digest solution was transferred to fresh eppendorf tubes. The gel pieces were extracted thrice with extraction buffer (1:1 ratio of ACN and 0.1% Triflouroacetic acid) and the supernatant was collected each time into the eppendorf above and then Speedvac till complete dryness. The dried pepmix was suspended in TA buffer.

### 2.12 Matrix-assisted laser desorption/ionization-time of flight mass spectrometry (MALDI-TOF/MS analysis)

The peptides obtained were mixed with HCCA matrix in 1:1 ratio and the resulting 2μl was spotted onto the MALDI plate [(MTP 384 ground steel (Bruker Daltonics, Germany)]. After air drying the sample, it was analyzed on the MALDI TOF/TOF ULTRAFLEX III instrument (Bruker Daltonics, Germany) and further analysis was done with FLEX ANALYSIS SOFTWARE (Version 3.3) for obtaining the PEPTIDE MASS FINGERPRINT. The masses obtained in the peptide mass fingerprint were submitted for Mascot search in “CONCERNED” database for identification of the protein.

### 2.13 Validation of Proteomic Results

#### 2.13.1 RNA isolation and cDNA synthesis

Total RNA was extracted from pre-metastatic (0 hrs APF) and post-metastatic (24 hrs APF) *Scrib* knockdown pupae using TRI reagent (TAKARA) as per manufacturer’s protocol. RNA extracted from age matched wild type pupae was used as control. The quantity of RNA was checked on a 1% agarose gel and documented using Gel Doc (BIO-RAD). High quantity RNA from both samples estimated by absorbance A260/280 ratio was reversibly transcribed to cDNA through reverse transcriptase PCR.

For cDNA preparation, 1µg of RNA, 1µl random hexamer (Applied Biosystems) and 11µl (0.1%) DEPC treated water was run for single cycle at 70^0^C for 5min. in a thermal cycler (BIO-RAD). To this 2µl 10mM dNTPs (Invitrogen) and 0.5µl (200U/µl) RNase inhibitor (Applied Biosystems) were added and run for single cycle at 25^0^C for 5 min. Finally, 1µl (100U/µl) reverse transcriptase (Ambion) was added and whole mixture was run at 37^0^C for 1h in a thermal cycler (Sure Cycler 8800, Agilent).

#### 2.13.2 Validation of newly identified candidates of *Scrib* knockdown induced metastasis Qualitative Real-time PCR (qRT-PCR)

Prepared cDNA of desired genotype was used as a template for quantification of differentially expressed genes identified by 2DE, normalized against GAPDH with specific probes Table.1 using SYBR (R) GREEN JUMPSTART TAQ Ready mix (Thermo Fisher Scientific) through Real-time thermal cycler analysis (ΔCT values).

**Table.1.**
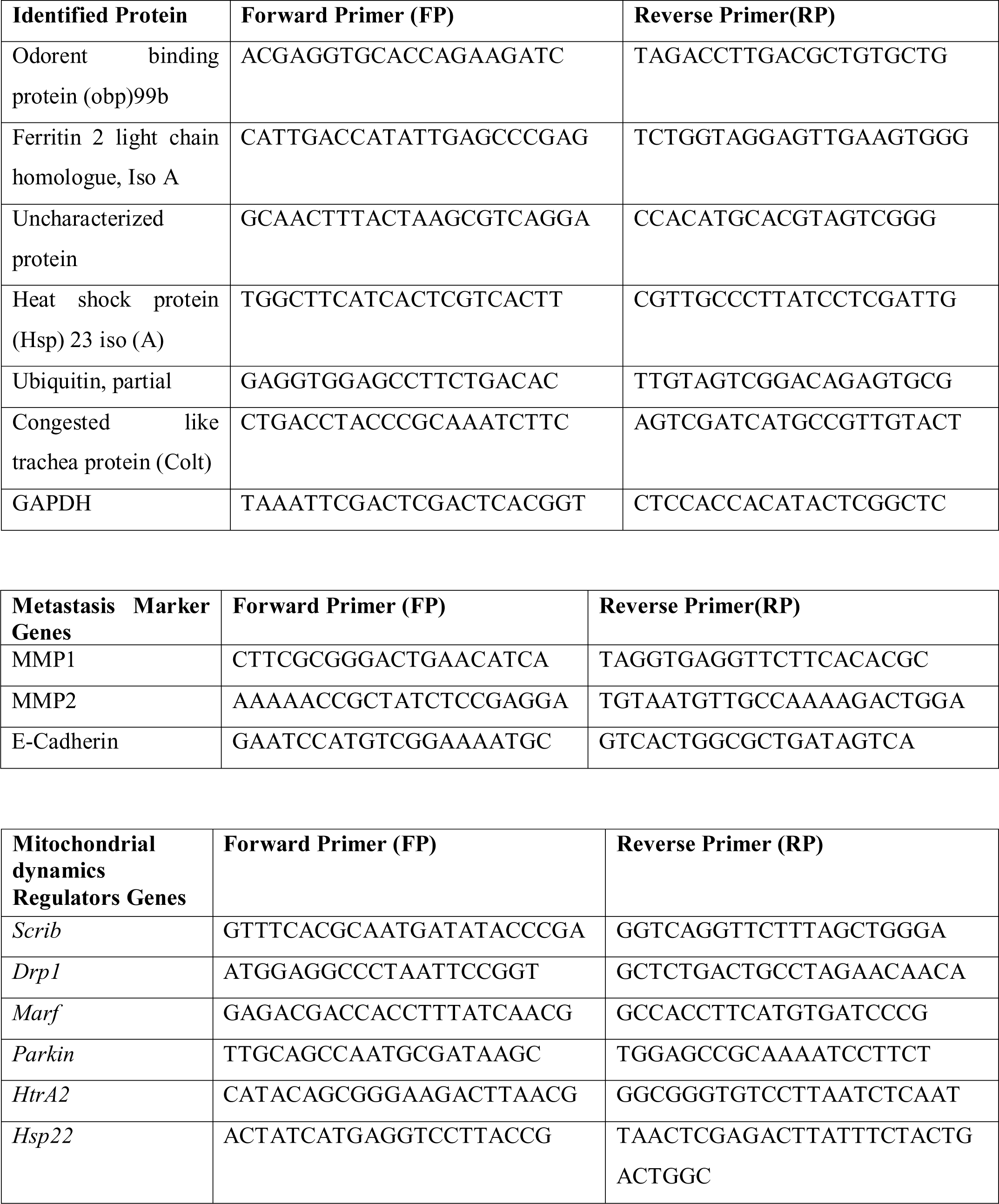
Specific primers (shown in 5’-3’ direction) for identified candidate genes.

#### 2.13.3 Western Blot

Total protein was extracted from pre-metastatic (0 hrs APF) and post-metastatic (24 hrs APF) *Scrib* knockdown pupae. Age matched wild type pupae were used as control. Bradford assay was used to measure the protein concentration. Equal amount of protein (100µg) were wet transferred after being separated by 10% SDS-PAGE onto PVDF membrane. Blot were probed with a cocktail of three MMP1primary antibodies (1:100, DSHB 3A6B4, 3B8D12 and 5H7B11), *Dlg* (1: 200, DSHB 4F3)and β-tubulin (E7). Proteins were detected using HRP linked anti-IgG secondary antibody and specific bands were detected using the ECL system. Gene expression was analyzed after normalizing β-tubulin expression using image J softwere.

#### 2.13.3 Metal content Analysis with Inductively Coupled Plasma (ICP)

For analysis of Fe^2+^ metal homeostasis, 15 pupae of wild-type and *Scrib* knockdown *Drosophila* pupae (24 hrs APF), were collected from culture bottles in glass petri plates and dried at 60^0^C for 12 hrs. Pupae were digested for analysis of total metal content in 5 ml 65% HNO_3_ (SDFCL, 20168 L05), boiled at 55-60^0^C for 30 min. Samples were cooled and filtered by Whatman paper and maintained to final volume upto 10 ml. The sample were analyzed against their respective standards of 0.5 ppm, 1 ppm and 1.5 ppm (Multi-element calibration standard 3 for ICP, 10µg/ml, PerkinElmer, L2-20MKBY1). The metal homeostasis analysis is done in Perkin-Elmer Optima 700 DV ICP optical emission spectrometer version 5.0. Data was analyzed based on the triplicate experiments.

##### Prussian Blue staining

Wing imaginal discs from third instar larvae of Oregon-R and UAS-*Scrib^RNA^*^i^/ Ptc-Gal4 were dissected in 1X PBS, fixed in 4% paraformaldehyde for 30 min, and permeabilizedwith 1% Tween 20 in 1X PBS for 15 min. To detect the ferrous iron in imaginal discs, the tissues were incubated in dark with freshly-made Prussian blue staining solution (2% K4[Fe(CN)6] and 2% HCL) for 1 hr, washed done 3 times with 1x PBS and mounted in 1X PBS and imaged on a Nikon microscope.

#### 2.13.4 Oxidative Stress Analysis

24hr pupae of *Scrib* knockdown and wild type strain were analyzed for H_2_O_2_ and Catalase assays in triplicates.

##### H_2_O_2_ (Hydrogen peroxide) Assay

The sample of desired genotype were homogenized in sodium phosphate buffer (50mM, pH 6.6) containing Na_2_HPO4 and NaH_2_PO4, and centrifuged 10,000rpm for 10 minute at 4°C. The supernatants were transferred to a fresh 1.5ml micro centrifuge tube and the concentration of protein is estimated by Bradford method. In 100µg sample protein was added to 1ml TiSO_4_ (0.1% in 20% H_2_SO_4_),and incubated for 10 minute at room temperature. The sample was centrifuged at 1000rpm for 20 minute at 4°C. Absorbance were taken at 410nm using multimode plate reader Synergy H1.

##### Catalase (CAT) Assay

The sample of desired genotype was homogenized in catalase extraction buffer (pH 8.0) containing Tris-HCl (50mM), EDTA (0.5mM) and 2% (w/v) PVP (polyvinyl pyrrolidone). The homogenized sample were centrifuged at 10,000rpm for 10 minute at 4°C. The supernatants were transferred to a fresh 1.5ml micro centrifuge tube and concentration of protein was estimated by Bradford method. The assay mixture contains 100µl crude total protein, 1ml potassium buffer (contain 100mM K_2_HPO4, 100mM KH_2_PO4) and 20mM of 8.8M H_2_O_2_ solution. The H_2_O_2_ consumption was measured 10times at 5s intervals and absorbance was recorded at 240nm. The catalase activity is calculated by subtracting highest value of absorbance to lowest value of absorbance 240nm divided by time log (5 sec) multiplied by extinction coefficient 0.036.

#### 2.13.5 *In silico* Analysis

Protein-Protein interaction signaling pathways were analyzed by STRING 9.0 web software. STRING is a functional protein connection networks analysis ( https://string-db.org) using protein accession number to analyze the connecting link between molecular signaling pathways and interaction associated between the differentially expressed proteins [19]. Further, Phylogenetic tree analysis was performed using latest version of Molecular Evolutionary Genetics Analysis (MEGA) 7.0.9 software [20] downloaded from www.megasoftware.net to determine evolutionary distance and genetic relationships [21]. The identified protein sequences downloaded from NCBI (National Center for Biotechnology Information) in FASTA format are identified by Gene Bank accession number. The sequences were aligned with MUSCLE using UPGMA method. The evolutionary distances were computed using the Poisson correction model [22].

#### 2.13.6 Statistical Analysis

Experiments were performed in triplicate and statically analyzed by Graph Pad Prism-5 offline software. All data were compared with one tailed student t-test or two-way ANOVA with Bonferroni posttests depending on experimental number of groups involved. Error bar denotes standard error of mean (SEM). (***p<0.001, **p<0.01 and ns p>0.05).

## 3. Results

### 3.1. Tracking metastasis from pre-metastatic phase to post-metastatic phase in *Scrib* knockdown tumor bearing pupae

To track metastasis in *Drosophila* cancer model, we used the GFP-tagged *Scrib* knockdown cancerous cells. GFP-labeled wild-type (WT) and *Scrib* knockdown pupae were analyzed for records of observation started from wandering larval stage wing imaginal discs to upto 84 hours, till complete adult wing formation where cuticle is transformed into a hard sealed puparium. GFP-labeled wild type pupae (UAS-GFP; Ptc-GAL4) used as control (Fig.1.A-L) were compared to tumor-bearing *Scrib* knockdown (*Scrib^RNAi^/*UAS-GFP;Ptc-GAL4) pupae (Fig.1.M-X). The third instar larvae crawling upward out of corn-meal agar medium and molt into a transparent white pre-pupa are consider as 0 hours pupae (Fig.1A and M); indicate pre-metastatic condition. GFP was observed in a single strip along the anterior-posterior (A/P) boundary of the 0 hours white transparent pupae in wing imaginal discs of control (Ptc-GAL4> UAS GFP), (Fig.1A), whereas in *Scrib* knockdown, a large undifferentiated mass of GFP-tagged cancerous cells were observed (Fig.1M). In control pupae, there is no apparent change in the expression pattern of GFP Till 12 hours (Fig.1C). Further, gradual development of GFP-tagged control wing discs into adult wings could be visualized in the thorax region (FIG.1D-L). In *Scrib* knockdown, the GFP-tagged cancerous cells were located in their original sites upto 12 hours as primary tumor (Fig.1O)(N=27;90%) but after 24 hours,(post-metastatic phase), cancerous cells dislodge from the primary tumors and colonize at distant sites of pupae, suggesting that *Scrib* knockdown tumor cells have the ability to get metastasized from primary sites to secondary sites (Fig.1P,n=22;73%).No visible changes were observed in the GFP pattern for the next 6hrs (Fig.1Q), but at 36hrs, further spreading of GFP-tagged mutant cells could be visualized (Fig.1R). Note marked dissemination of fluorescent cells from anterior toward posterior end of pupae at 48 hours (Fig.1S) which persisted upto 84 hours (Fig.1T-X). Percentage of pupae exhibiting metastasis upon loss of *Scrib* function is shown in Fig.1.Y.

**Fig.1.**
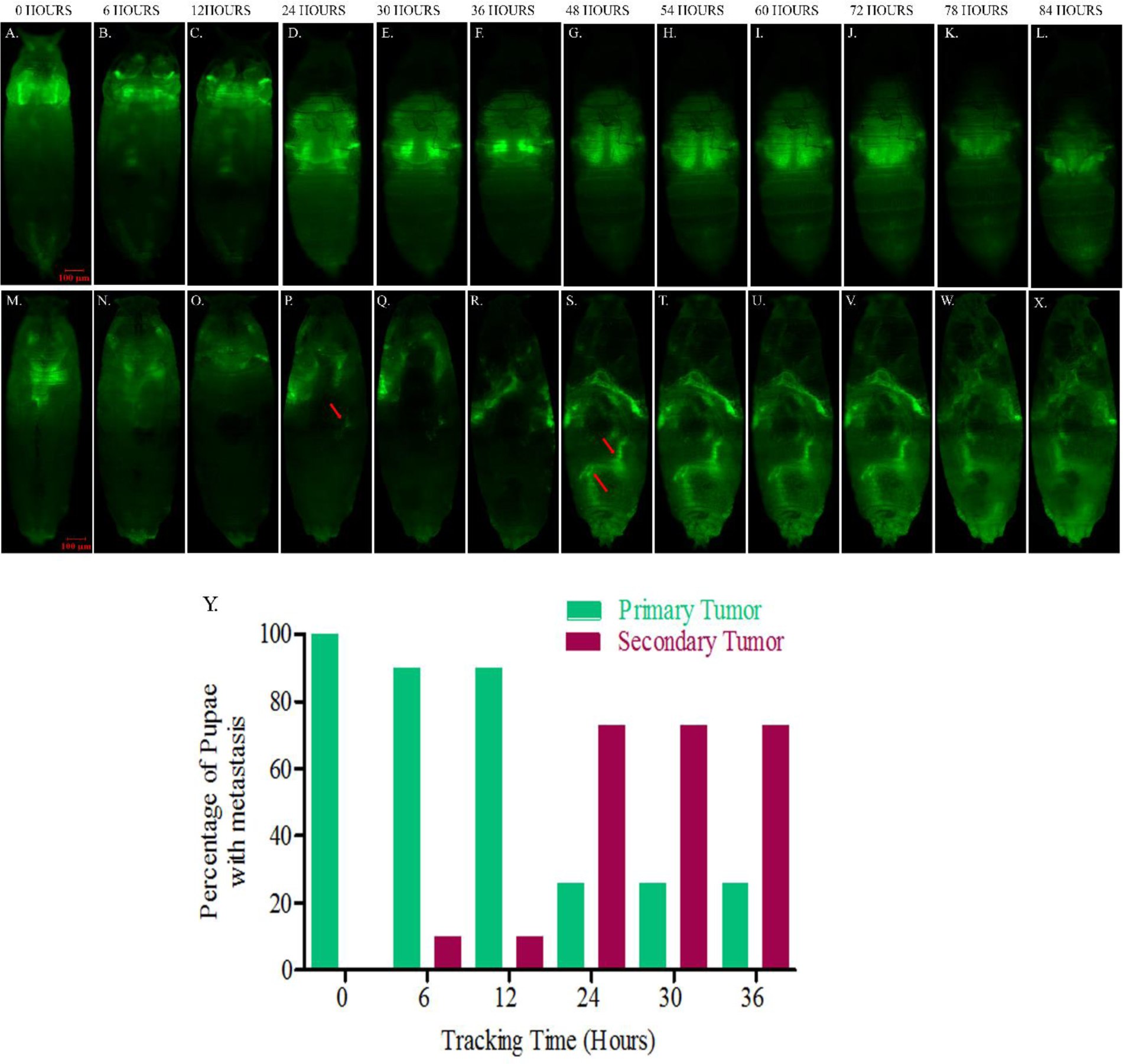
Tumor progression and metastasis phenotypes in *Drosophila melanogaster* were scored about 84 hours after puparium formation (APF). Scale bars: 100µM. In WT pupae GFP expression is in restricted regions (A-L). *Scrib* knockdown mutant cells develop into primary tumors but not secondary tumors development upto 12 hours APF (M-O). Secondary tumor (red arrow) get metastasized from primary site appear after 24 hours APF (P-R) during post-metastatic condition. Note the markedly movement of GFP-tagged cells at 48 hours (S) and remain constant up to 84 hours (T-X). Y.Metastasis tracking chart represents the percentage of pupae with metastasis upon loss of *Scrib* at different time points noted upto 84 Hours but since no changes occur in pupae after 36 hours, above histogram of percentage of pupae with metastasis is marked up to 36 hours where Green bars show percentage of primary tumors and red bars show secondary tumors.

### 3.1. Metastatic ability of *Scrib* tumor cells depends on MMP1 expression in *Drosophila* pupae

We examined MMP1 expression in *Scrib^RNAi^* compared to wild-type during 0hrs APF, pre-metastatic (24hrs APF), and post-metastatic phase of *Drosophila* pupae. We performed real-time PCR and western blot for detection of MMP1 mRNA and protein expression respectively (Fig.2). The transcript levels of MMP1 during pre-metastatic condition (0 hours APF) in *Scrib* knockdown pupae were 0.6 fold downregulated, however during post-metastatic condition (24 hours APF) were 4.7 fold upregulated when compared to wild-type(Fig.2.B). Further, the transcript levels of MMP2 in *Scrib* knockdown pupae were 0.4 fold downregulated in pre-metastatic condition, however during post-metastatic condition they were 1.2 fold upregulated when compared to wild-type (Fig.2.C).However, the transcript levels of E-cadherin in *Scrib* knockdown pupae were not significantly different in comparison to wild type(0.7 fold and 0.8 fold downregulated between pre-metastatic and post-metastatic condition, respectively)(Fig.2. D). The band intensity clearly indicates significant down regulation of MMP1 protein during 0hrs APF pre-metastatic phase and significantly upregulation during 24hrs APF post-metastatic phase compare to wild-type pupae (Fig.2. E and F). Immunostaining for MMP1 antibody reveal higher expression of MMP1 protein in *Scrib* knockdown tumor as compared to wild-type (Fig.3).

**Fig.2.**
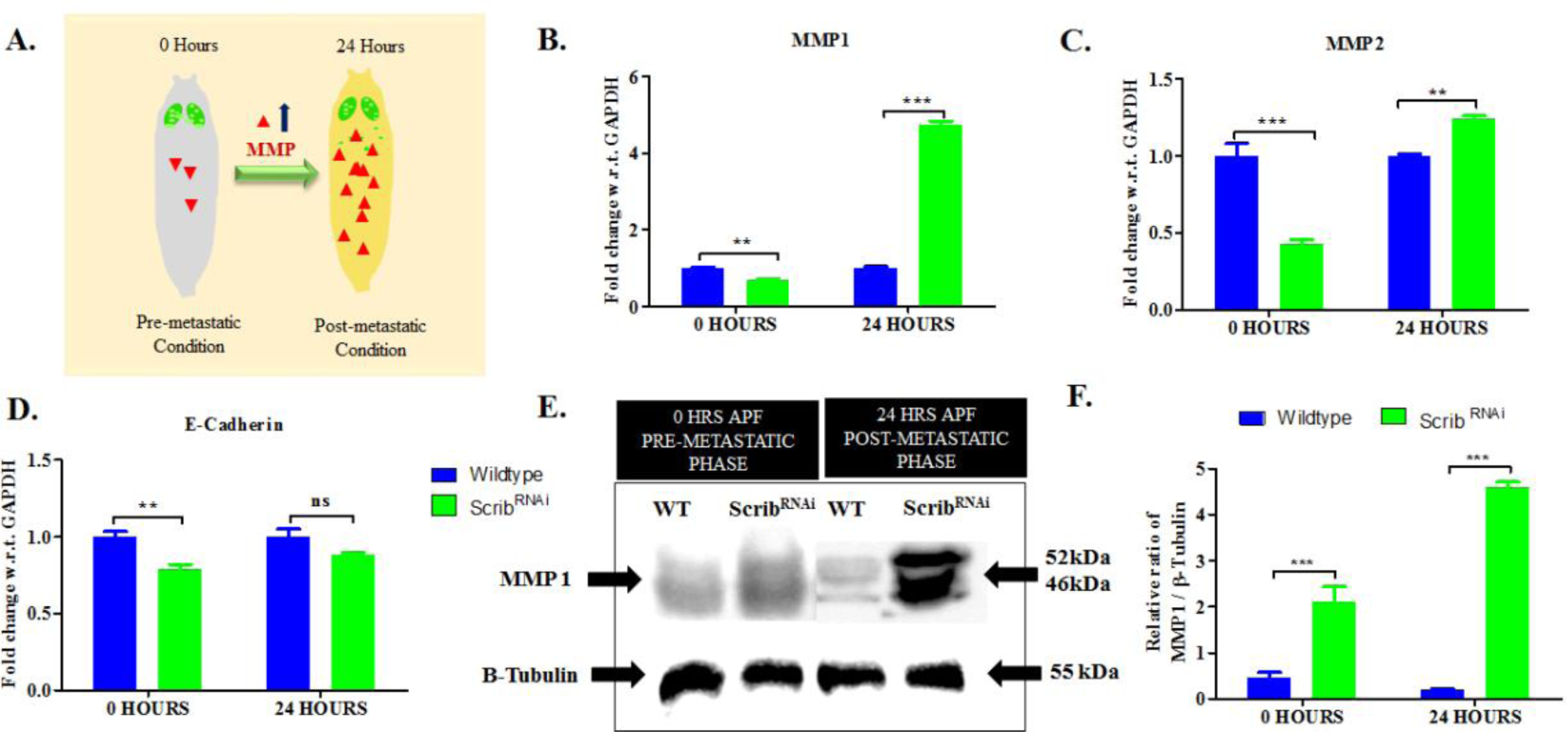
Relative mRNA and protein expression level of MMPs in *Scrib* knockdown *Drosophila* pupae. A. Schematic diagram representing level of MMP protein during 0 hours and 24 hours APF in pre-metastatic and post-metastatic condition respectively of *Drosophila* pupae. Histogram representative (B, C and D) of RT-PCR was performed using primers against GAPDH, MMP1, MMP2 and E-Cadherin during pre-metastatic and post-metastatic condition of *Drosophila* pupae. Western blot analysis of MMP1 (E) and the relative ratio of band intensity (F) revealed higher level of MMP1 as compared to wild-type during post-metastatic condition of *Drosophila* pupae. An equal amount of total protein was loaded, as verified with anti-β tubulin. Analysis of data was done using two-way ANOVA with Bonferroni posttests. (***p<0.001, **p<0.01 and ns p>0.05).

**Fig.3.**
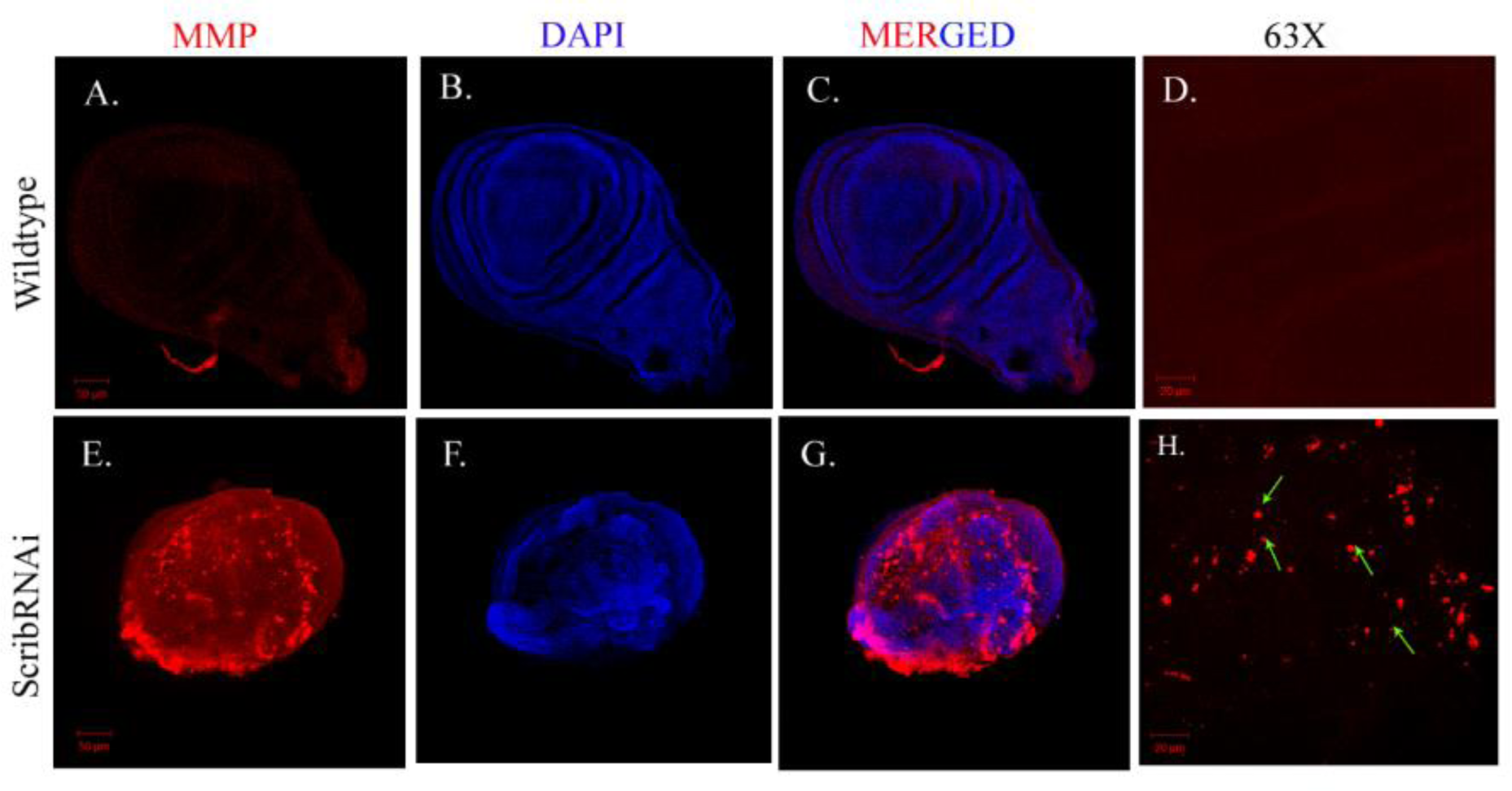
Confocal projection showing MMP1 Immunostaining. (A-D) MMP1 expression is not detectable in wild-type (red) in comparison to *Scrib* knockdown tumorous cells (E-H). D and H represent 63X magnified images of Wild-type with low MMP1and *Scrib^RNAi^* with overexpression of MMP1 (indicated by green arrow), respectively.

### 3.2. Identification of Differentially expressed proteins in post-metastatic pupae

MALDI-TOF/MS analysis was used to examine differential protein expression profile after 24 hrs of pupae formation during post-metastatic condition of *Scrib* knockdown pupae verses wild-type. The protein spots on the 2-D gel electrophoresis were visualized and matched by difference in spot intensities between wild-type and *Scrib^RNAi^* mutant cancerous tissues (Fig.4). The expression of 18 spots automatically detected from gel were analyzed with PD Quest software basic 8.0.1, BIO-RAD. Of those, 10 protein spots appeared to have higher protein intensity levels and 8 protein spots appeared to have lower levels in cancerous tissue than Wild-type. Six most significantly altered protein spots, of which three were upregulated (Fig.5.A-C) and three downregulated (Fig.5.D-F) in cancerous tissue compared to wild-type were chosen for MALDI-TOF/MS analysis. Details of novel proteins identified from MALDI-TOF/MS analysis is given in Table 1. The upregulated proteins include Odorant-binding protein 99b (DmelObp 99b), Ferritin 2 light chain homologue, isoform A (Dmel Fer2LCH) and CG13492, an uncharacterized gene/protein). On the other hand, downregulated proteins include Heat shock protein 23 (Dmel Hsp23), Ubiquitin, Partial (DmelRpl40) and Congested-like trachea protein (Dmel Colt).

**Fig.4.**
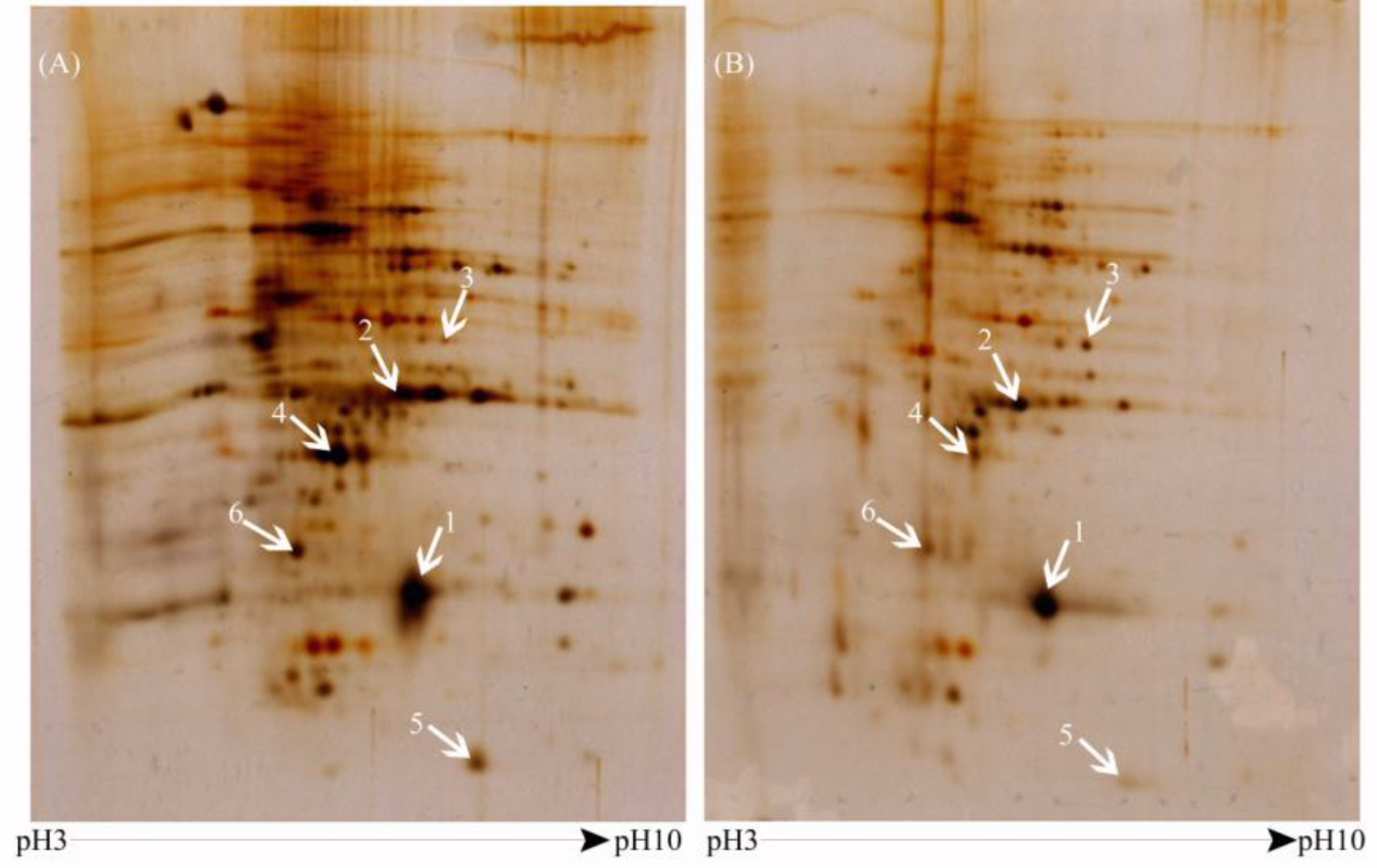
Two-dimensional gel electrophoresis (2-DE) images of wild type (A) and *Scrib* knockdown pupae (B). The six differentially expressed protein spots as specified by numbers 1-6 indicated by white arrows were picked for MALDI-TOF MS analysis. The pH range (pH3-pH7) of BIO-RAD IPG (immobilized pH gradients) strip is indicated in the below of figure.

**Fig 5.**
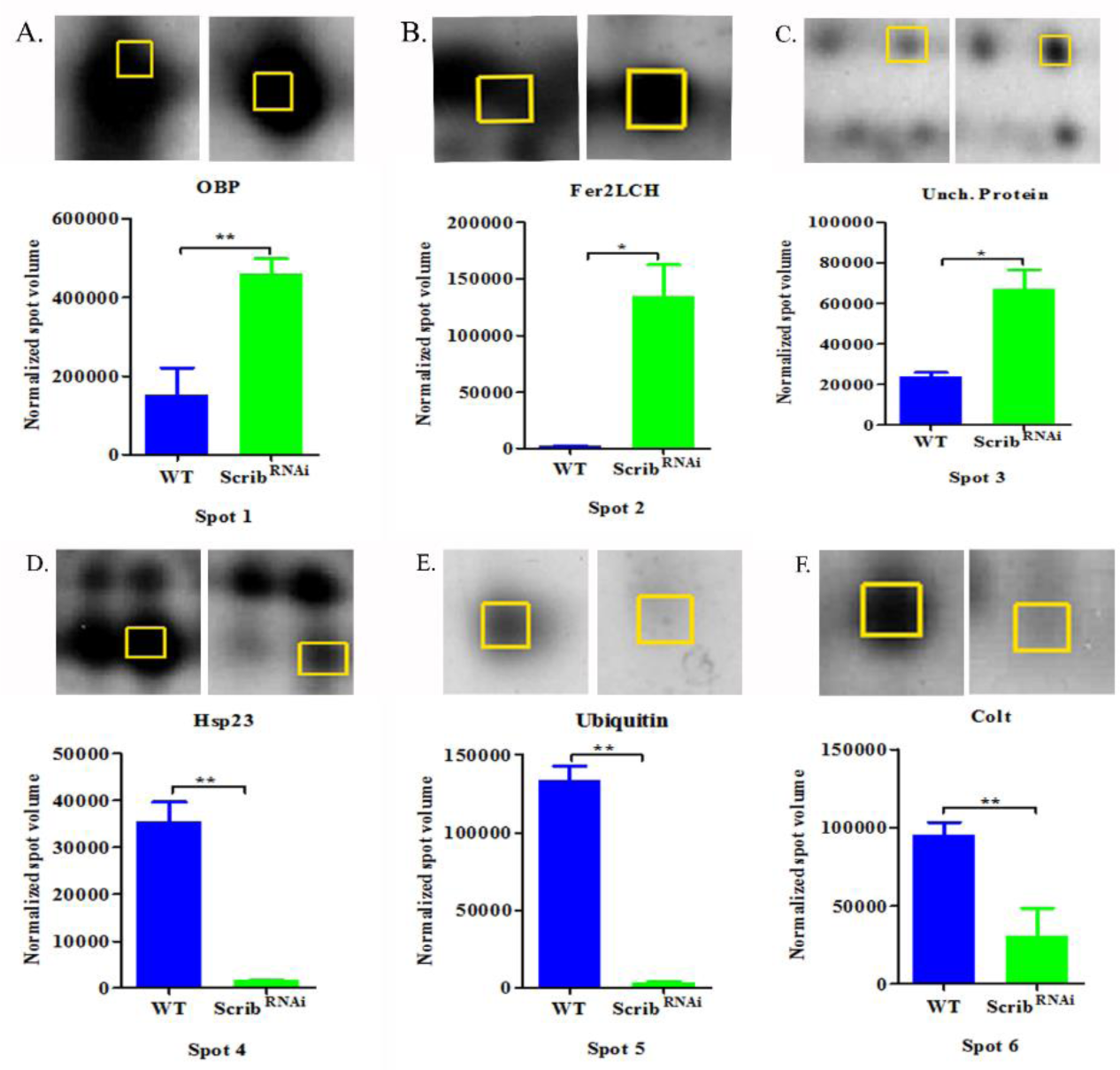
Histogram represents the normalized protein intensity values of each spot in wild-type compared with *Scrib^RNAi^* and magnified image shown above the graphs. The protein spots 1-3 are significantly up-regulated(A-C) and the protein spots 4-6 are down-regulated (D-F) in *Scrib* knockdown cells as compared to wild-type. Analysis of data was done using t tests (p<0.05).

### 3.3. Validation of Differentially expressed proteins during pre-metastatic and post-metastatic phases of *Scrib^RNAi^*

Real-time PCR analysis was performed to estimate the mRNA transcript levels of newly identified *Scrib* associated genes during pre and post-metastatic condition of tumor bearing pupae at 0hrs and 24hrs APF respectively verses wild-type pupae. The transcript levels of OBP99b at 0 hr and 24hr APF were marginally but not significantly altered between wild type and ***Scrib^RNAi^*** transcript levels in pre and post metastatic *Scrib* knockdown pupae were 0.98 fold down-regulated, and 1.2 fold upregulated, respectively, when compared to that wild-type, (Fig.6.A). The transcript levels of Fer2LCH protein during pre-and post-metastatic condition in *Scrib* knockdown pupae were 1.35fold and 1.10 fold up-regulated, respectively, when compared to that wild-type (Fig.6.B). The transcript levels of CG13492 during pre-and post-metastatic condition in *Scrib* knockdown pupae were surprisingly 0.40 fold and 0.33 fold down-regulated, respectively, compared to wild-type (Fig.6.C), rather than being upregulated. The transcript levels of Hsp23 protein during pre-and post-metastatic condition in *Scrib* knockdown pupae were significantly up-regulated (2.26 fold and 3.15 fold, respectively),compared to wild-type (Fig.6.D). The transcript levels of Ubiquitin during pre-metastatic condition (0 hours APF) in *Scrib* knockdown pupae were 1.59 fold significantly upregulated, however during post-metastatic condition (24 hours APF) significantly, they were 0.46 fold down-regulated compared to wild-type (Fig.6.E). Similarly, the transcript levels of Colt protein during pre-metastatic condition (0 hours APF) in *Scrib* knockdown pupae were 2.61 fold upregulated, however during post-metastatic condition (24 hours APF) they were notably down-regulated by 0.58 fold, when compared to that wild-type (Fig.6.F).

**Fig.6.**
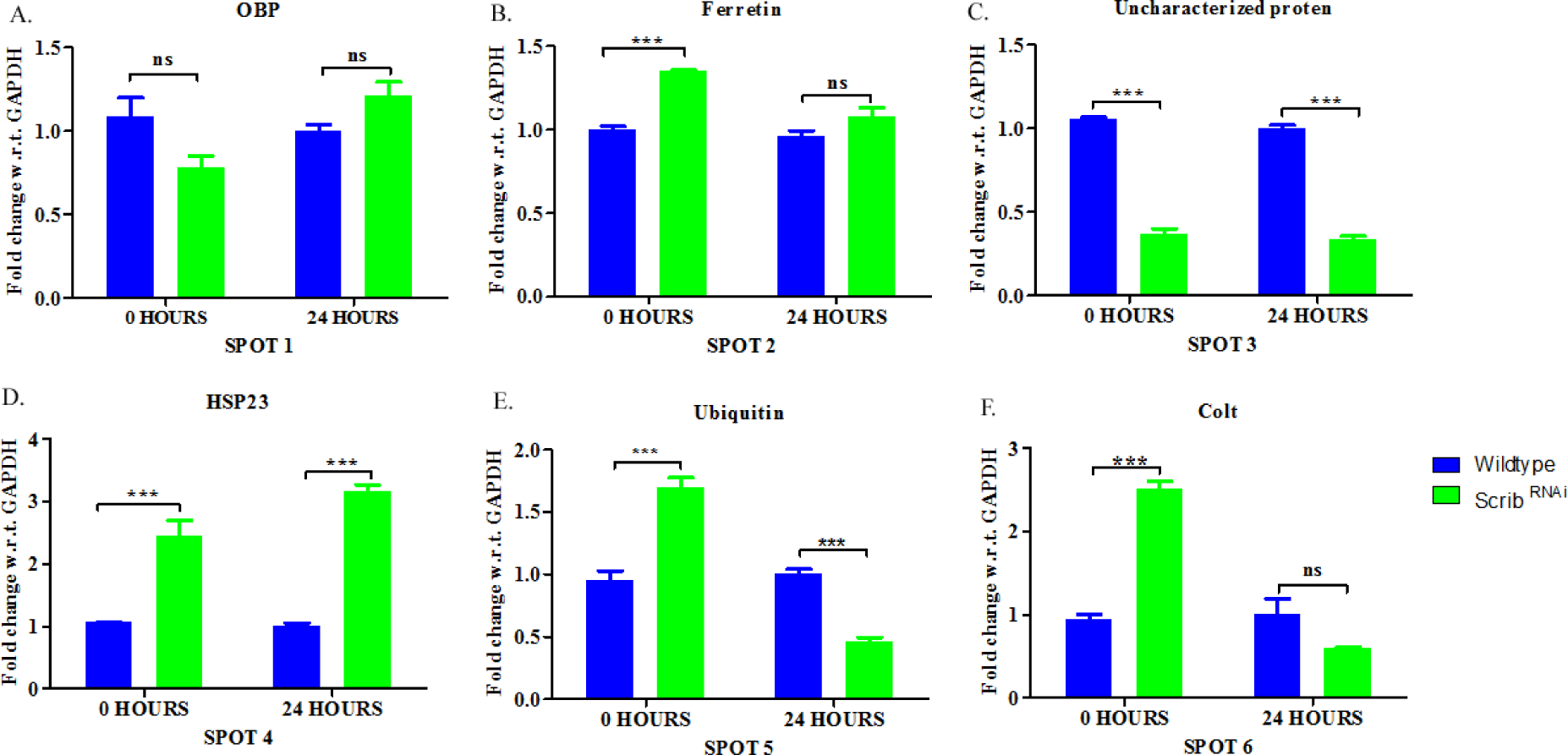
Relative mRNA expression of genes corresponding to the identified proteins, as measured by quantitative real time PCR. RT-PCR was performed using primers against GAPDH and newly identified *Scrib* associated genes, OBP, Ferritin, Uncharacterized protein (CG13492), HSP23, Ubiquitin and Colt (A-F). The data was normalized to GAPDH mRNA level and fold change in mRNA level of identified genes is shown. Analysis of data was done using two-way ANOVA with Bonferroni posttests. (***p<0.001, and ns p>0.05).

### 3.4. *Scrib* knockdown tumorous cells exhibit altered Iron Homeostasis

The novel *Scrib* associated protein identified, ferritin, an iron storage protein shows higher expression in *Scrib* knockdown tumor bearing pupae as compared to WT. To further validate the increase in iron production in tumorous pupae, we performed ICP and Prussian Blue staining. The ICP results showed slightly enhanced level of Fe^2+^ in *Scrib* knockdown cells compared to wild-type (Fig. 7.A). Likewise, Prussian Blue staining also revealed that Ferrous ion levels were increased in tumorous pupae as compared to WT (Fig.7.B).

**Fig.7.**
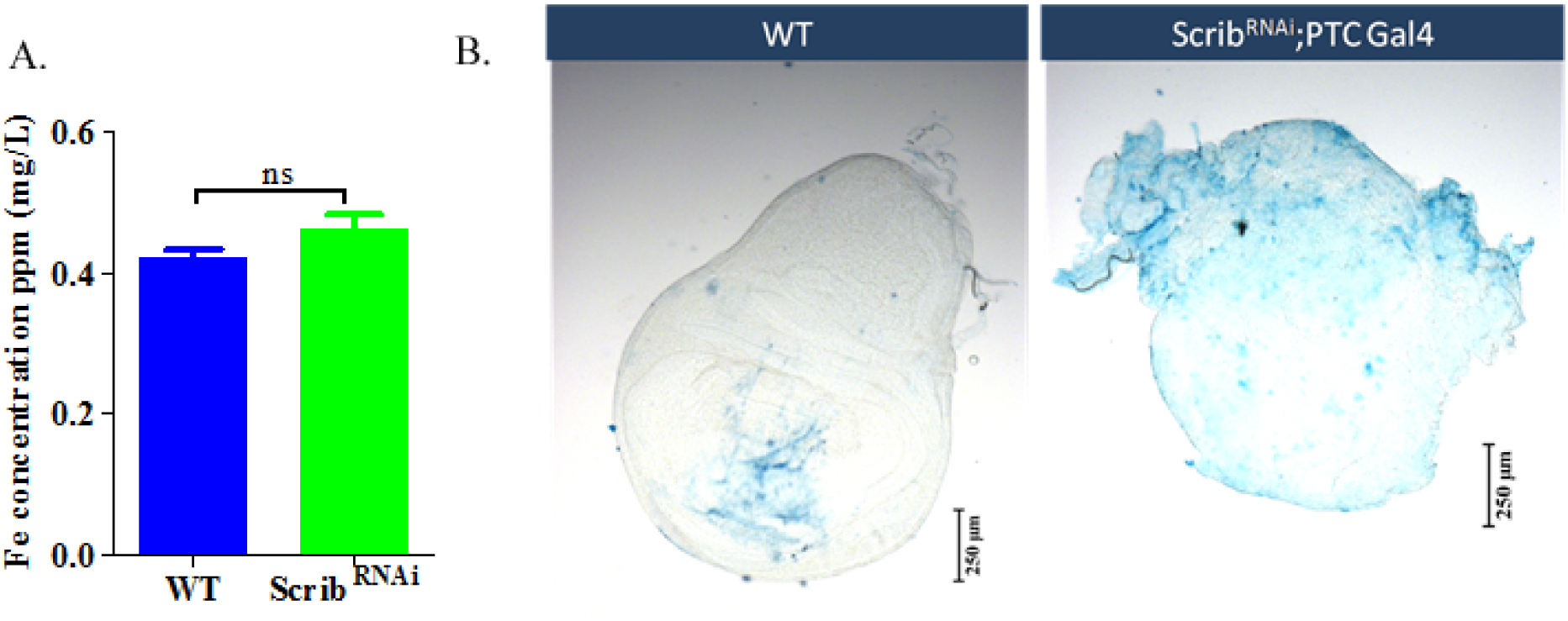
Altered Fe homeostasis in *Scrib* knockdown pupae compared to wild-type. The Fe^2+^ levels,examined by ICP were marginally enhanced in *Scrib* knockdown pupae as compared to Wild-type (A), Analysis of data was carried out using t tests (ns p>0.05). Prussian blue staining of wing imaginal discs isolated from WT and *Scrib* knockdown larvae (B). Note the increase in Fe2+ levels in *Scrib^RNAi^* as compared to wild type.

### 3.5. Loss of *Scrib* disrupted mitochondrial dynamics and elevated oxidative stress during post-metastatic condition of *Drosophila* pupae

The relative mRNA expression of mitochondrial dynamics regulators*Drp1, Marf, HtrA1, Parkin* and *Hsp22* were assessed during post-metastatic phase (APF 24 hrs) of *Scrib* knockdown and WT pupae using Real time PCR. The Drp1 and Hsp22 mRNA transcript level were marginally upregulated, Marf and *Parkin* mRNA levels were slightly down regulated, whereas*HtrA1*transcripts were significantly down regulated during post-metastatic condition of *Scrib* knockdown tumorous pupae as compared to wild-type pupae (Fig. 8.A). Further oxidative stress marker catalase showed slightly lower levels as compared to wild type (Fig.8.B), whileH_2_O_2_levels were significantly enhanced (Fig.8.C) suggesting an enhanced ROS production during post-metastatic phase of *Scrib* knockdown compared to wild-type.

**Fig.8.**
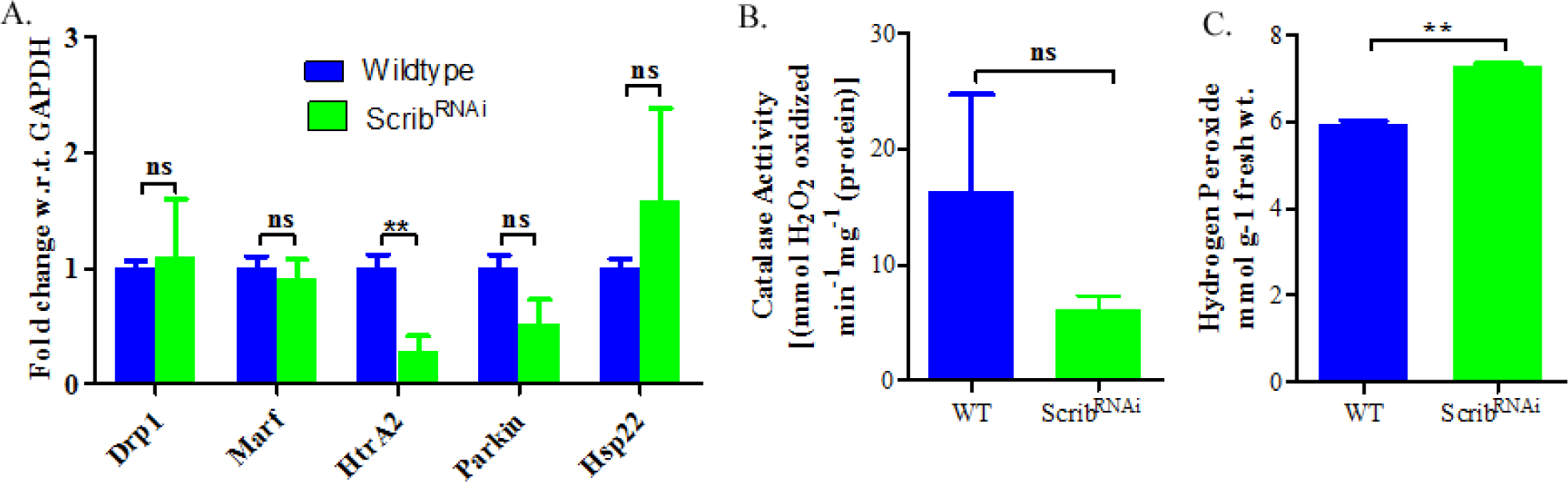
Alteration in mitochondrial dynamics regulator genes during post-metastatic condition 24hrs APF of *Scrib* knockdown pupae analyzed by Real-time PCR. Histogram represents the mRNA transcript level of mitochondrial dynamics regulator genes in *Scrib* knockdown pupae compared to Wild-type (A). The catalase antioxidant enzyme activity and H_2_O_2_ production level measured in *Scrib* knockdown pupae compared to wild-type pupae (B and C).Analysis of data was done using t tests (ns p>0.05, ** p<0.05).

### 3.6. In silico Analysis for protein-protein interaction network of newly identified proteins

Using STRING database, a protein-protein interaction network diagram was automatically generated for 6 newly identified *scrib* interacting proteins. The protein RpL40 for Ubiquitin, partial is located at the central region of protein-protein network has an experimental link to colt, scribble and molecular chaperone, Hsp23. The RpL40 has functional partners RpL15, RpL18A, RpS28b, RpLP2 and RpS19a. All functional partners have co-expression with RpL40. The RpS28b is a gene neighbor of RpL40. Fer2LCH is co-expressed with RpL15 and RpL18A, whereas RpL18A interacts with Fer2LCH. Fer2LCH, Hsp23, Scribble and colt are located in the outer region of a generated network of protein-protein interaction diagram. The differentially expressed proteins Obp99b and uncharacterized protein (CG13492) did not show any interaction with other differentially expressed proteins in the protein-protein network as shown below in the Fig.9.

**Fig.9.**
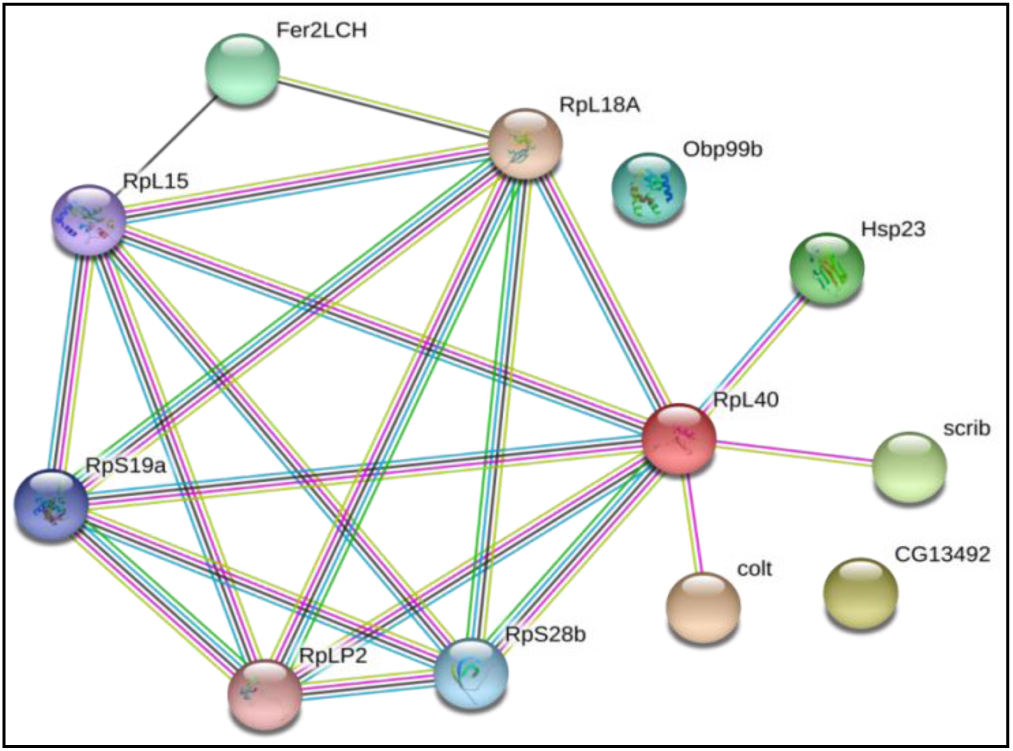
The protein-protein interaction network diagram as analyzed by STRING software 9.0 web software. 5 different colored lines represent 5 types of connections needed to predict the association of *Scrib* with differentially expressed proteins. This network contains 12 nodes and 20 edges to show protein-protein interactions. Sky blue line for curated databases, purple line for experimental determined association, green for gene neighborhood connection and black for co-expression predicted interaction for differentially expressed genes.

### 3.7. Phylogenetic tree analysis

In-silico evolutionary analysis was used for comparison of protein sequences at the molecular level. The optimal tree with the sum of branch length 6.43383242 is generated by MEGA 7 software (Fig.10). The Phylogenetic analysis involved seven amino acid sequences. The similar protein sequences lie very close evolutionary relationship. In Phylogenetic tree, Ubiquitin, Partial with Congested-like trachea protein, Ferritin 2 light chain homologue, isoform A with Heat shock protein 23 share a common ancestor and Uncharacterized protein Dmoj_GI21508 with Scrib and they both share common evolutionary relationship with Odorant binding protein. Thus branching point in constructed evolutionary Phylogenetic tree shows how two protein amino acid sequences are closely related with different proteins.

**Fig.10.**
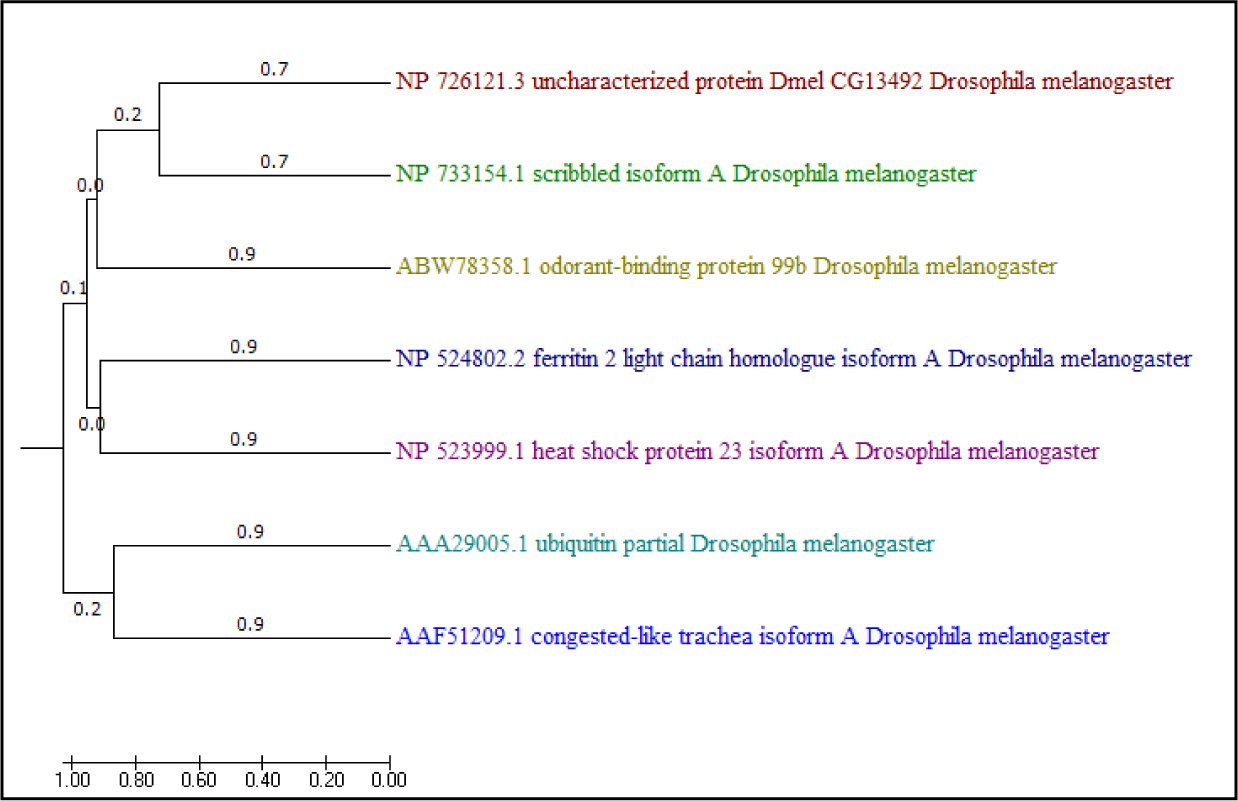
The Phylogenetic tree was constructed using MEGA 7.0.9 software. The gene bank accession number for protein sequences used are listed as follows, Obp99b (ABW78358.1), Fer2LCH (NP_524802.2), uncharacterized protein, CG13492 protein (NP_726121.3), Hsp23 (NP_523999.1), Ubiquitin, Partial (AAA29005.1), Colt (AAF_51209.1) Scrib (NP_733154.1). The FASTA sequences were aligned with MUSCLE method and analyzed by UPGMA method. The scale indicates branch lengths of evolutionary distances used in Phylogenetic tree construction.

## Discussion

The major cause of more than 90% cancer associated deaths is due to metastasis and also the main reason of failure of cancer treatment [23]. Tumor metastasis is the process by which an uncontrolled malignant growth of primary tumor cells migrates from its origin to the new distant sites where secondary tumors form, but exact molecular and cellular signaling involved in the cascade of metastatic events remains elusive. The big challenges for cancer researchers are the identification of novel metastasis-associated proteins marker targets, which might add a major contribution to the study of biological aspects of metastasis cancer. To explore this, here, we have generated a system for the study of metastatic cancer in *Drosophila* pupae through knockdown of *Scrib* in wing imaginal disc using UAS-GAL4 bipartite approach and for first time we performed timeframe analysis for tracking metastasis events from pre to post-metastasis phase of *Drosophila* pupae development with the help of GFP reporter (n=30). The *Scrib* knockdown in the wing imaginal disc resulted in primary tumor development, which metastasized throughout the pupa, as revealed by GFP reporter protein. *Drosophila* larvae bearing malignant tumor progressed normally throughout larval development but never properly pupated as wild-type pupae (Fig.1). Till the 12 hours, early pupae carried primary tumors (Fig.1O) in wing imaginal discs (N=27;90%) but after 24 hours during post-metastatic phase, GFP tagged cancerous cells floating in the hemolymph, suggests that *Scrib* knockdown tumor cells get metastasized from primary sites to secondary sites (n=22, 73%). This study revealed that the majority of primary tumors spread from primary site to other parts of pupae due to excessive cell proliferation and migration of *Scrib* knockdown cells between 12-24 hours as revealed by appearance of GFP-tagged cells to other parts of pupae that may be because of enhance in MMP1 secretion(Fig.1P). Further, we explore the role of MMP1 which is sufficient for encouraging secondary tumor formation and metastasis upon loss of *Scrib* from pre-metastatic phase to post-metastatic phase APF of *Drosophila* Pupae.MMP is important player involved in metastasis and maintenance of pupal development. We found that MMP1 expression was upregulated during 24 hrs APF, compared to wild type as revealed by RT-PCR, western blotting as well as immunostaining (Fig.2.B, Fig. 2.E and F and Fig.3, respectively). We also checked MMP2 expression and found 1.2 fold upregulation during 24hrs APF in post-metastatic phase compared to 0hrs APF during pre-metastatic phase verses wild-type (Fig.2.C). Further, E-Cadherin transcript levels, were downregulated during pre-metastatic phase (0 hrs APF) and post-metastatic phase (24hrs APF) compared to wild-type pupae (Fig.2.D). Our findings revealed that metastasis biomarker,MMP1, MMP2 and E-Cadherin are involved in regulation of metastasis upon loss of *Scrib* alone, without cooperative involvement of any other oncogenes. S*crib* knockdown alone in epithelial cells is sufficient for malignant tumor growth in *Drosophila* pupae.We find that the S*crib^RNAi^* tumor bearing pupae show100% lethality after 36 hours.

Further, to find out key players involved in tumor progression and metastasis, we utilized proteomic differential display analysis using two-dimensional gel electrophoresis (2-DE) and Matrix assisted laser desorption/ionization-time of flight (MALDI-TOF) mass spectrometry (MS). Proteomics is a systematic research technology used to identify novel proteins involve in tumor progression and metastasis [24]. We identified 6 differentially expressed novel metastasis associated proteins during post-metastatic condition (24hrs APF) in *Scrib* knockdown tumor bearing pupae. The upregulated proteins include Odorant-binding protein 99b (DmelObp 99b), Ferritin 2 light chain homologue, isoform A (Dmel Fer2LCH) and Uncharacterized protein (Dmel CG13492). On the other hand, downregulated protein includes Heat shock protein 23 (Dmel Hsp23), Ubiquitin, Partial (DmelRpl40) and Congested-like trachea protein (Dmel Colt) (Fig. 4 and Fig. 5). Further, mRNA expression level of these novel proteins were checked during 0hrs APF in pre-metastatic phase and 24hrs APF in post-metastatic phase of *Drosophila* pupae using real-time PCR (Fig. 6). We found, Fer2LCH, Hsp23, Ubiquitin and colt were up-regulated and Obp 99b and Uncharacterized protein-CG13492 were down-regulated at 0hrs during pre-metastasis condition whereas, at 24hrs during post-metastasis condition Obp 99b, Fer2LCH and Hsp23 were up-regulated and uncharacterized protein-CG13492, Ubiquitin and colt gene expression were down-regulated in *Scrib* knockdown tumor bearing pupae as compare to WT pupae.

As results show, the fer2LCH mRNA transcript level were shown significant upregulation during pre and post-metastatic phase in *Scrib* knockdown pupae development. To further validate the same, we performed ICP (Inductively coupled plasma) and Prussian blue staining, which confirmed the upregulation of Fe^2+^ levels in *Scrib* knockdown cells compared to wild-type (Fig. 7). Further, we also measured the level of an antioxidant catalase enzyme and H_2_O_2_ production in *Scrib* knockdown cells compared with WT pupae. We found that low level of catalase activity and higher production of H_2_O_2_ support the excess ROS production. Thus our studies revealed that at 24 hrs APF during post-metastatic phase, the novel metastasis associated ferritin protein supporting ROS production to stimulate the cancer cells proliferation and metastasis in *Scrib* knockdown cells (Fig. 8).

Furthermore, in-silico studies of these novel protein using protein-protein interaction analysis by STRING software shows ferritin protein has interaction with Hsp23 protein and phylogenetic tree analysis using MEGA software also shows common ancestor of ferritin and Hsp23 protein (Fig. 9 and Fig. 10). According to previous reports, the inverse correlation of Hsp23 protein expression and gene transcripts shows decline in Hsp23 protein level and maximum detection of Hsp23 transcripts up to 50 hours of pupae formation is due to low stability of Hsp23 protein [25,26,27]. In our study, we also quantifying protein and mRNA expression level after 24 hours of pupae formation during post-metastatic condition of tumor bearing pupae and found high accumulation of Hsp23 mRNA and a decline in protein expression. Overexpression of sHsps has also been reported to protect mammalian cells from oxidative stress [28]. In this study, we also found an elevated level of Hsp23 during pre-metastatic and post-metastatic phase at 0 hrs and 24 hrs APF respectively to protect cancer cells from the excess ROS production. Small Hsps help to prevent protein aggregation or target damaged proteins towards two molecular pathways, the Ubiquitin-proteasome system (UPS) and the Autophagy-lysosome degradation system [29]. In this context, we identified Ubiquitin protein upon *Scrib* knockdown in cancerous pupae through proteomic analysis.

Ubiquitin, Partial (DmelRpl40, spot 5) is a small protein that is covalently attached to target protein for degradation by Ubiquitin-proteasome system [30]. Ubiquitin molecule is encoded by four different encoded genes, UBA52, RPS27a, Ubiquitin B (UBB) and Ubiquitin C (UBC) genes that are highly homologous in eukaryotes. [31]. UBA52 and RPS27a genes code for a single copy of ubiquitin fused to ribosomal protein L40 (RpL40) and ribosomal protein S27a (Rp S27a) respectively. The Ubiquitination controls cellular processes of tumorigenesis, like cell-cycle progression and apoptosis and also play important role in removal of harmful proteins [32]. In this study, we also found an elevated level of Hsp23 with upregulation of ubiquitin during pre-metastatic condition 0hrs APF and downregulation of Ubiquitin during post-metastatic condition upon loss of *Scrib* showing connective link between them.

Further, among downregulated proteins, congested-like trachea protein (Dmel Colt, spot 6) belongs to the mitochondrial carrier family. The colt gene of *Drosophila* regulates gas-filling of tracheal system and cell proliferation during larval stage to generate single-layered epithelium of wing imaginal discs [33]. In our study, colt gene expression is downregulated during post-metastatic condition at 24hrs APF so, from these we hypothesized that downregulation of the colt gene expression could affect the mitochondrial dynamics. To further confirmed that colt gene is related to the mitochondrial dysfunction, we checked the status of mitochondrial dynamics regulators genes, Drp1, Marf, HtrA2 and *Parkin* expression in *Scrib* knockdown tumorous pupae during post metastasis phase by qRT-PCR. Also we checked the mRNAs transcript level of mitochondrial localized small heat shock protein, Hsp22 and found that upregulation of *Drp1 and Hsp22* gene whereas, downregulation of *Marf, HtrA2, Parkin* gene in *Scrib* knockdown cells (Fig.10.A). Therefore, study suggested that the novel involvement of *Colt* proteinduring post-metastatic phase at 24hrs APF could affect the mitochondrial dynamics in *Scrib* knockdown tumorous tissues trigger cancer cell metastasis.

Recently, Song et al., 2023identified novel role of *Apt* gene, which also regulate tracheal system and knockdown of *Apt* gene in background of *Scrib^RNAi^* enhance MMP1 secretion through activation of JNK pathway trigger cancer metastasis in *Drosophila*[34]. Similarly, Colt mitochondrial associated proteins also regulating *Drosophila* tracheogenesis could affect the JNK signaling pathway and may have role in cancer metastasis. Colt protein have not been previously reported in regulation of cancer cell proliferation and we first time report novel role of colt protein in regulation of *Scrib* knockdown cancer cell metastasis events during post-metastasis phase after 24 hours of pupae formation in *Drosophila*.

Thus, our observation suggesting that upregulation of ferritin protein and down regulation of colt protein expression might trigger the MMP1 secretion and excessive ROS production which leads to disruption of mitochondrial dynamics upon loss of *Scrib* stimulate cancer cell proliferation and metastasis. The altered expression of this novel protein required further investigation in the direction of mitochondrial dysfunction associated with loss of *Scrib* in prevention of cancer cell metastasis.

## Conclusion

The finding of our present study that *Scrib* alone, without cooperative association of *Ras* or other oncogenes, has potential to metastasized to distant parts of pupae during a particular timeframe of the pupal development. Further, we found that MMP1 expression is increased during 24hrs post-metastatic condition in *Scrib* knockdown pupa compared to wild type. The current studies suggest that the increased production of MMP1 in *Scrib* knockdown wing imaginal tissues facilitates their metastasis events in tumor bearing pupae. A comparative proteomic analysis of metastatic pupae resulted in newly identified novel proteins, ferritin and colt proteins, during cancer cell migration and metastasis. The systematic tracking of malignant tumor cells from an early stage of pupae to post metastasized stage, when primary tumor metastasized to distant sites, provides good understanding on the involvement of various novel proteins and their specific role towards better therapeutic interventions of tumor progression and metastasis. Further research on these newly identified proteins in the context of Scrib will result in the treatment of cancer.

## Acknowledgments

This work was supported by a research grant (P07/598) awarded to SS by the Science and Engineering Research Board, India and ICMR-Senior Research Fellowship (ICMR-SRF) to JS. Authors are highly grateful to IoE (Institution of Eminence) for financial support under incentive grant and SATHI for central facility. The authors would like to greatly acknowledge ISLS-BHU and UPE-BHU for Nanodrop, Real time PCR and other routine facilities.

## Supplementary Information

### TUMOR GENERATION

**Fig.S.1.**
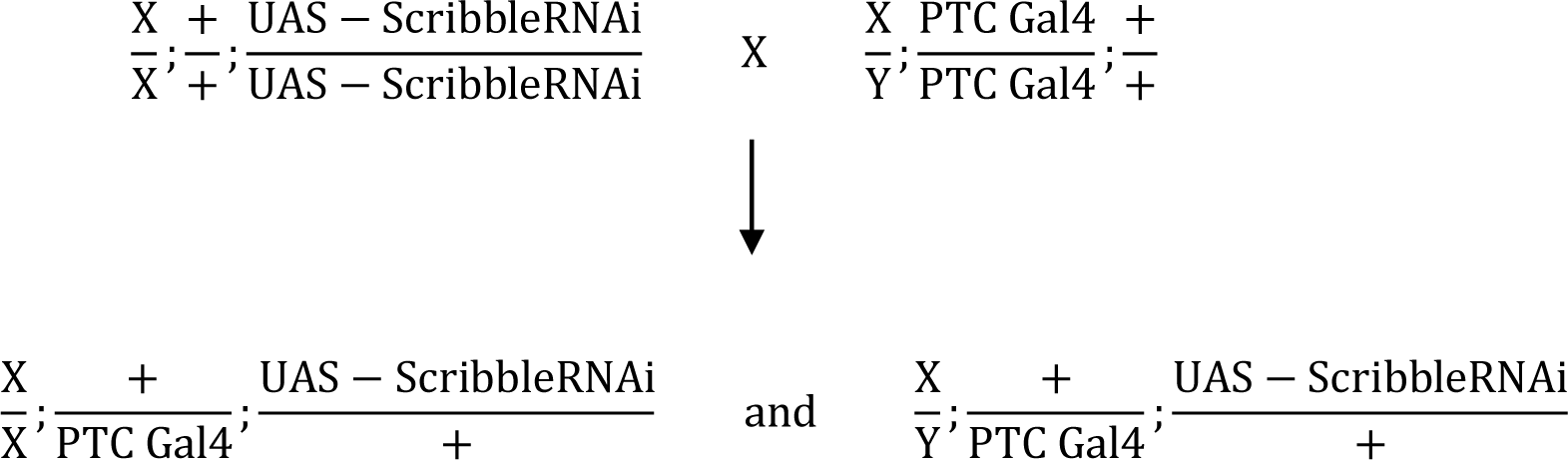
Schematic representation for knockdown *Scribble* specifically along the A-P boundary of *Drosophila* wing disc. 15 virgin females flies of UAS-*scribble*^*RNAi*^ cross set with 15 males flies of PTC-Gal4 in vial at 24^0^C.

### Scheme for making PTC Gal4 as GFP

**Fig.S.2.**
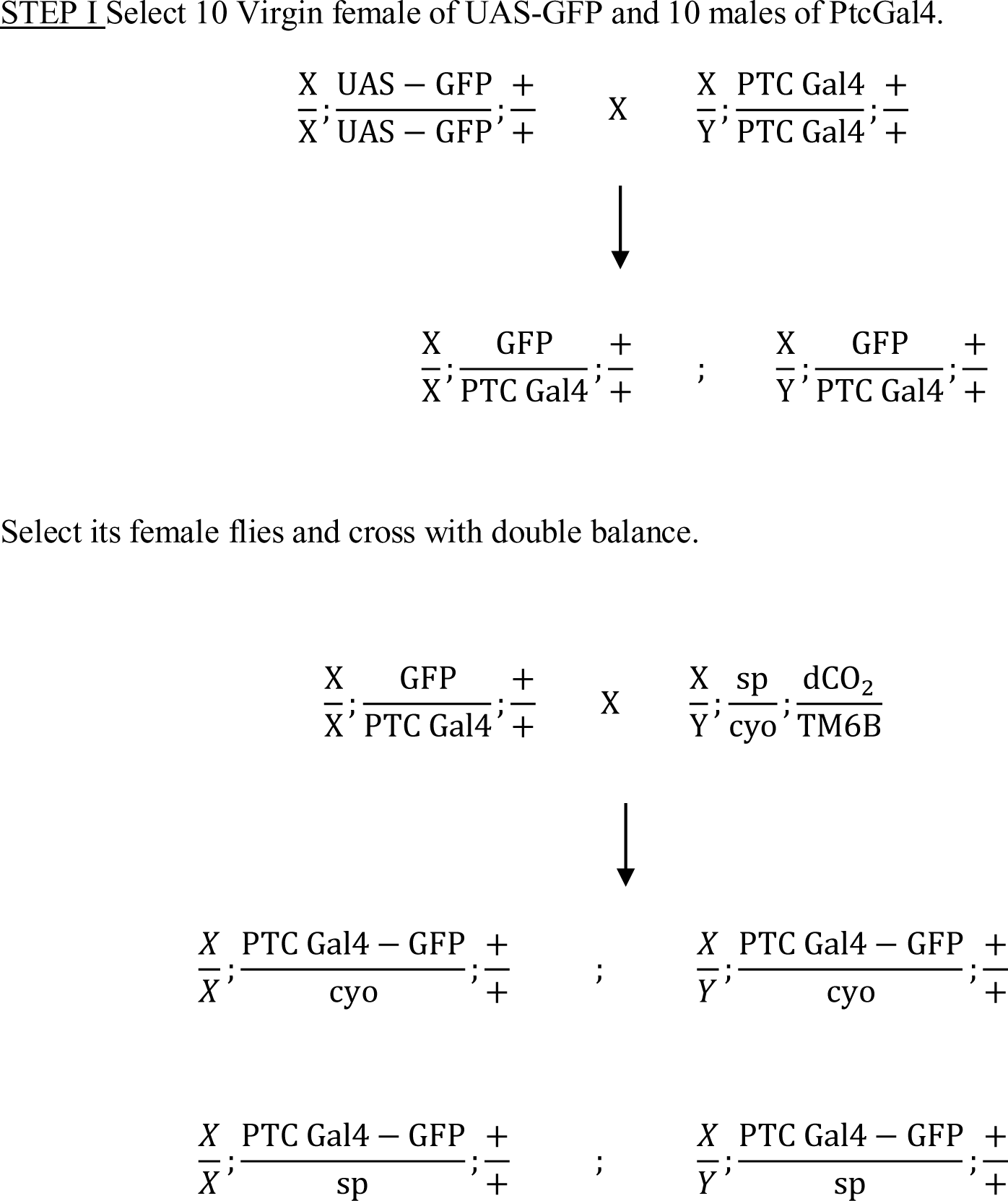
To monitored tumor progression and metastatic behavior in living transparent pupa, genetically label these wing discs specific PTC-GAL4 with a visible marker such as green fluorescent protein (GFP).

